# *Artemisia annua* hot-water extracts show potent activity *in vitro* against Covid-19 variants including delta

**DOI:** 10.1101/2021.09.08.459260

**Authors:** M.S. Nair, Y. Huang, D.A. Fidock, M.J. Towler, P.J. Weathers

## Abstract

**Ethnopharmacological relevance:** For millennia in Southeast Asia, *Artemisia annua* L. was used to treat “fever”. This medicinal plant is effective against numerous infectious microbial and viral diseases and is used by many global communities as a source of artemisinin derivatives that are first-line drugs to treat malaria.

**Aim of the Study:** The SARS-CoV-2 (Covid-19) global pandemic has killed millions and evolved numerous variants, with delta being the most transmissible to date and causing break-through infections of vaccinated individuals. We further queried the efficacy of *A. annua* cultivars against new variants.

**Materials and Methods:** Using Vero E6 cells, we measured anti-SARS-CoV-2 activity of dried-leaf hot-water *A. annua* extracts of four cultivars, A3, BUR, MED, and SAM, to determine their efficacy against five fully infectious variants of the virus: alpha (B.1.1.7), beta (B.1.351), gamma (P.1), delta (B.1.617.2), and kappa (B.1.617.1).

**Results:** In addition to being effective against the original wild type WA1, *A. annua* cultivars A3, BUR, MED and SAM were also potent against all five variants. IC_50_ and IC_90_ values based on measured artemisinin content ranged from 0.3-8.4 μM and 1.4-25.0 μM, respectively. The IC_50_ and IC_90_ values based on dried leaf weight (DW) used to make the tea infusions ranged from 11.0-67.7 μg DW and 59.5-160.6 μg DW, respectively. Cell toxicity was insignificant at a leaf dry weight of ≤50 μg in the extract of any cultivar.

**Conclusions:** Results suggest that oral consumption of *A. annua* hot-water extracts (tea infusions), could provide a cost-effective therapy to help stave off the rapid global spread of these variants, buying time for broader implementation of vaccines.

## 1.0 Introduction

The global SARS-CoV-2 (Covid-19) pandemic has infected ~220 million people and killed >4.5 million (https://coronavirus.jhu.edu/map.html). Numerous variants have rapidly evolved (https://www.who.int/en/activities/tracking-SARS-CoV-2-variants/). The delta variant is currently the most transmissible to date (RO = 5-7) (Nunes-Vaz and Macintyre, 2021), causing break-through infections in vaccinated individuals (Gupta et al., 2021). Effective and approved small molecule-based therapeutics are still lacking. Recently, we showed that hot-water extracts of dried leaves of seven cultivars of the medicinal plant, *Artemisia annua* L., used for millennia to treat malaria fever (Hsu, 2006) and sourced from four continents, prevented SARS-CoV-2 replication *in vitro* (Nair et al., 2021). Recently, anti-SARS-CoV-2 efficacy of *A. annua* extracts was independently confirmed (Zhou et al., 2021).

Antiviral efficacy inversely correlated to artemisinin (ART) content (Nair et al., 2021). Others also observed that compared to *A. annua*, *A. afra*, a related perennial species lacking ART, was equally effective vs. SARS-CoV-2 with IC_50_ values of 0.9-3.4 and 0.65 mg/mL extract, respectively (Nie et al., 2021). Although these results indicated that both *A. annua* and *A. afra* have potent anti-SARS-CoV-2 activity *in vitro* and that the effect is not ART dependent, it was unclear whether *A. annua* is effective against emerging variants.

Here we report *in vitro* efficacy against new variants of four of the seven originally studied *A. annua* cultivars.

## 2.0 Methods and Materials

### 2.1 Plant material, extract preparations, and artemisinin analyses

Hot-water extracts (tea infusions) were prepared from dried leaves of four cultivars of *Artemisia annua* L. (SAM, MASS 00317314; BUR, LG0019527; A3, Anamed; MED, KL/015/6407) and analyzed as detailed in (Nair et al., 2021). Briefly, hot-water extracts were made from 10 g dried leaves/L that were boiled in water for 10 min, sieved to remove solids, and filter-sterilized (0.22 μm) prior to storage at −20°C. For ART analysis, tea infusions were extracted and analyzed by gas chromatography-mass spectrometry as detailed in (Martini et al., 2020). ART contents in μg/mL were: 42.5 for A3; 20.1 for BUR; 59.4 for MED4; and 149.4 for SAM (Nair et al., 2021).

### 2.2 Viral culture and infection

Vero E6 (ATCC CRL-1586) cell cultivation and viral infection were detailed in (Nair et al., 2021). SARS-CoV-2 isolates (USA WA1; alpha, B1.1.7; beta, B1.351; gamma, P.1; delta, B.1.617.2; kappa, B.1.617.1) were sourced from BEI Resources (www.beiresources.org). Viruses were titrated upon propagation to determine their tissue culture infectious dose (TCID) in Vero E6 cells, aliquoted and frozen at −80°C until further use. Multiplicity of infection (MOI) was 0.1 as documented by (Liu et al., 2020).

### 2.3 Assays for determining drug inhibition of SARS-CoV-2 and cell viability

Extract dilutions were incubated for 1 h in wells of 96-well tissue culture plates containing a monolayer of Vero E6 cells seeded the prior day at 20,000 cells/well. Extracts were added at equal ART amounts based on the content of each extract. One hour later SARS-CoV-2 virus was added to each well at a final MOI of 0.1. Cells were cultured for 3 days at 37oC in 5% CO_2_ and were scored for cytopathic effects as previously detailed (Liu et al., 2020). Results were converted into percent of control. Drug concentrations were log transformed. Concentration of drug(s) that inhibited virus by 50% (*i.e.*, IC_50_), and concentration of drug(s) that killed 50% of cells (*i.e.*, CC_50_), were determined via nonlinear logistic regressions of log(inhibitor) versus response-variable dose-response functions (four parameters) constrained to a zero-bottom asymptote by statistical analysis. Viability of Vero E6 cells post extract treatment was already reported in Nair et al. (2021) for the same extracts.

### 2.4 Chemicals and reagents

Reagents were from Sigma-Aldrich (St. Louis, MO). EMEM (Cat # 30-2003) and XTT reagent (Cat # 30-1011k) were from ATCC. Renilla-Glo was from Promega (E2720).

### 2.5 Statistical analyses

All *in vitro* anti-SARS-CoV-2 analyses were done at least in triplicate. Plant extract analyses had n≥6 independent assays. IC_50_ and IC_90_ values were calculated using GraphPad Prism V8.0.

## 3.0 Theory/Calculation

*A. annua* hot-water infusions already shown to be effective against SARS-CoV-2 also are effective against newly emerging variants.

## 4.0 Results and Discussion

*A. annua* hot-water extracts inhibited the recently evolved variants of SARS-Cov-2 (Figure 1) with calculated IC_50_ values normalized to the ART content of each tea infusion ranging from 1.1 - 7. 9 μM for the gamma, delta, and kappa variants. Although already reported by (Nair et al., 2021), WT(WA1), alpha, and beta variants were included for direct experimental comparison (Figure 1; Table 1). The lowest IC_50_ values were from the BUR cultivar and the highest were from the SAM cultivar. As previously shown (Nair et al. 2021), there was an inverse correlation between ART in extracts and anti-viral efficacy. The lowest ART content (BUR) yielded the greatest potency (the lower the IC_50_, the more potent the drug/extract), providing evidence that ART is not the only active antiviral agent in these extracts. The Nie et al. (2021) preprint further validated that ART was not the only anti-SARS-CoV-2 compound in the extracts by showing that aqueous extracts of the PAR cultivar of *Artemisia afra*, an *Artemisia* species lacking ART, had an IC_50_ of 4.1 mg/mL, within the range of 3.1-13.0 mg dried extract/mL of the *A. annua* cultivars studied therein. As already reported for extracts used in this study, no cytotoxicity was observed at a dry weight of ≤50 μg in the extract of any cultivar (Nair et al. 2021).

**Figure 1.**
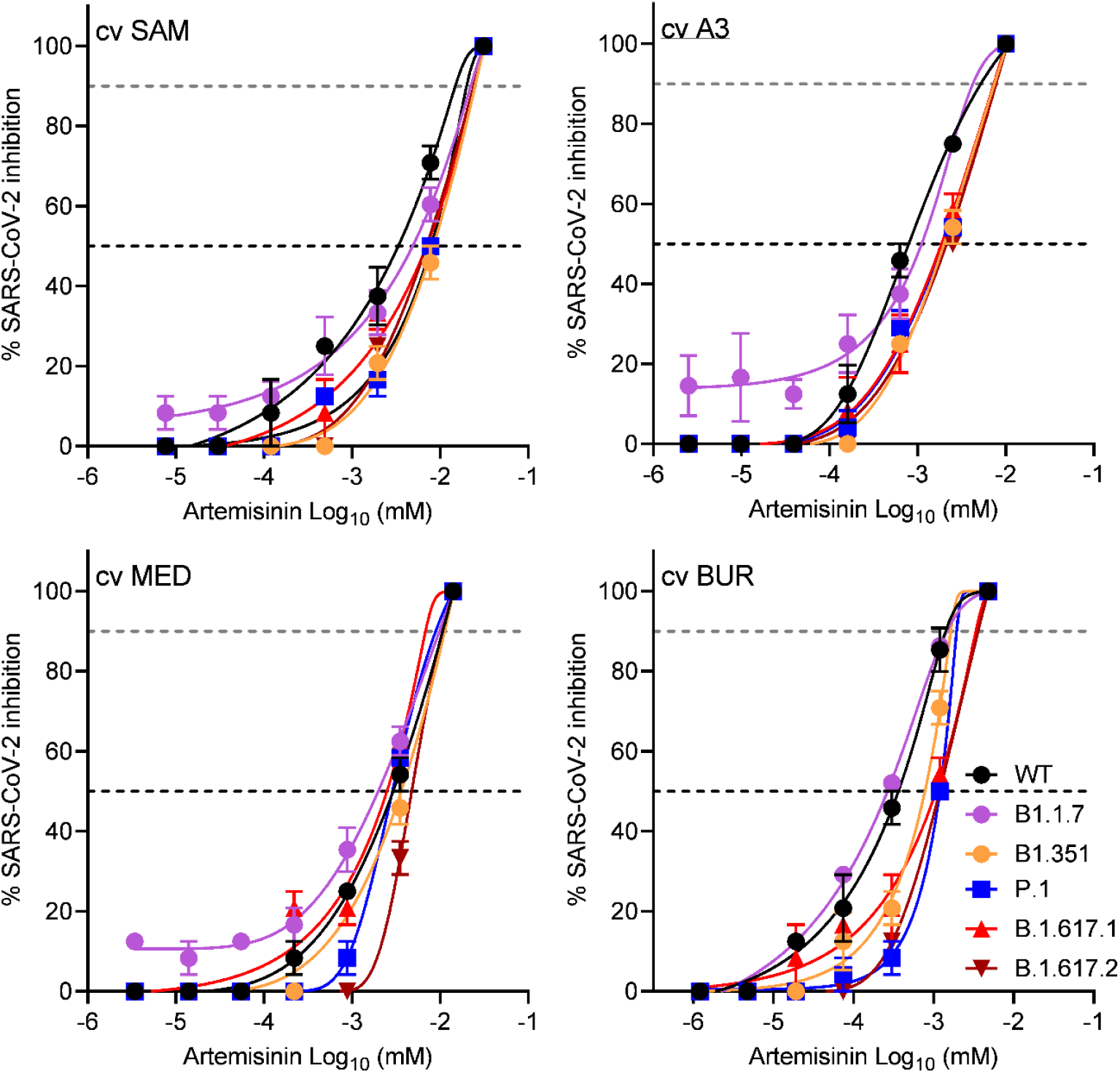
SARS-CoV-2 variant inhibition by four cultivars of *A. annua* hot water extracts normalized to their artemisinin content and compared to WT. WT, USA/WA1; variants: B.1.1.7, alpha; B.1.351, beta; P.1, gamma; B.1.617.1, kappa; B.1.617.2 delta) at a multiplicity of infection (MOI) of 0.1 in Vero E6 cells. Data for alpha and beta variants are extrapolated from Nair et al. (2021). Data are plotted from an average of three replicates ± SE.

**Table 1.** Potency of *A. annua* hot-water extracts (10g/L) against 6 strains of SARS-CoV-2 based on either artemisinin content or leaf dry weight (DW).

| Cultivar | Potency normalized to artemisinin content (μM) |  |  |  |  |  |  |  |  |  |  |  |
| --- | --- | --- | --- | --- | --- | --- | --- | --- | --- | --- | --- | --- |
|  | IC<sub>50</sub> μM artemisinin |  |  |  |  |  | IC<sub>90</sub> μM artemisinin |  |  |  |  |  |
|  | <u>WA1*</u> | <u>B.1.1.7*</u> | <u>B.1.351*</u> | <u>P.1</u> | <u>B.1.617.1</u> | <u>B.1.617.2</u> | <u>WA1*</u> | <u>B.1.1.7*</u> | <u>B.1.351*</u> | <u>P.1</u> | <u>B.1.617.1</u> | <u>B.1.617.2</u> |
| SAM | 3.4 | 4.9 | 8.4 | 7.9 | 7.0 | 7.0 | 14.9 | 22.3 | 25.0 | 20.0 | 24.8 | 24.1 |
| A3 | 0.8 | 1.1 | 2.0 | 1.9 | 1.9 | 2.1 | 5.2 | 4.2 | 7.4 | 6.5 | 7.3 | 7.8 |
| BUR | 2.9 | 2.0 | 3.6 | 2.9 | 2.5 | 4.8 | 10.7 | 9.8 | 11.3 | 8.9 | 6.8 | 10.6 |
| MED | 0.4 | 0.3 | 0.8 | 1.2 | 1.1 | 1.2 | 1.4 | 1.5 | 1.6 | 1.9 | 3.4 | 3.6 |
|  | Potency normalized to dry mass of leaves used in tea infusion (μg) |  |  |  |  |  |  |  |  |  |  |  |
|  | IC<sub>50</sub> μg leaf DW |  |  |  |  |  | IC<sub>90</sub> μg leaf DW |  |  |  |  |  |
|  | <u>WA1</u> | <u>B.1.1.7</u> | <u>B.1.351</u> | <u>P.1</u> | <u>B.1.617.1</u> | <u>B.1.617.2</u> | <u>WA1</u> | <u>B.1.1.7</u> | <u>B.1.351</u> | <u>P.1</u> | <u>B.1.617.1</u> | <u>B.1.617.2</u> |
| SAM | 21.5 | 31.3 | 53.7 | 50.7 | 45.0 | 45.1 | 95.6 | 143.1 | 160.4 | 128.0 | 159.3 | 154.7 |
| A3 | 15.7 | 22.1 | 39.6 | 38.2 | 37.0 | 42.4 | 103.7 | 84.1 | 147.9 | 150.2 | 145.3 | 154.7 |
| BUR | 15.1 | 11.0 | 32.5 | 50.1 | 44.7 | 49.8 | 59.5 | 61.2 | 68.6 | 80.4 | 143.3 | 150.4 |
| MED | 41.7 | 28.2 | 51.5 | 41.0 | 37.0 | 67.7 | 152.5 | 139.7 | 160.6 | 127.0 | 145.3 | 151.6 |

\* Values taken from Nair et al. (2021).
IC<sub>50</sub> and IC<sub>90</sub> are values where virus is 50% and 90% inhibited. Data are an average of three replicates.

Although Zhou et al. (2021) also showed that *A. annua* hot-water extracts had anti-SARS-CoV-2 efficacy, it is difficult to compare their IC_50_ values because they did not test the same viral strain, use the same plant cultivars, or make their extracts and apply them to viral infected cells using the same procedures. Using *A. annua* aqueous extracts against the BavPat 1/2020 strain of SARS-CoV-2 in Vero E6 cells, the IC_50_ values were 390 and 260 μg dried extract/mL for pretreated and treated cells, respectively. For pretreatment, extract was added 1.5 h before virus infection and for treatment, drug was added 1 h after virus infection. Ethanolic extracts yielded IC_50_ values about 50% lower, and thus were more potent than the aqueous extracts. To compare results of both studies, we calculated the dry mass of leaves equivalent to their reported IC_50_ to be 941.2 mg. The IC_50_ mass reported in (Nair et al., 2021) ranged from 13.5-57.4 μg, varying by cultivar. The leaf dry mass IC_50_s in this study for the gamma, delta, and kappa variants ranged from 38.2-50.7, 42.4-67.7, and 37.0-45.0 μg leaf DW, respectively. The three orders of magnitude difference between this study and Zhou et al. (2021) likely result from the above noted differences in methodology.

ART and its derivatives have some anti-SARS-CoV-2 activity (Cao et al., 2020; Gendrot et al., 2020a; Gendrot et al., 2020b; Nair et al., 2021; Zhou et al., 2021). However, in those reports where there are direct comparisons with *Artemisia* extracts, ART is not the only active phytochemical, suggesting there are other antiviral compounds in the plant. *A. annua* contains a rich assortment of identified phytochemicals (Ferreira et al., 2010), some of which have activity against human coronavirus proteins. For example, quercetin and myricetin have inhibitory activities against SARS-CoV NTPase/helicase with IC_50_s of 0.1 and 2.7 μM, respectively, and luteolin has an IC_50_ of 10.6 μM against SARS-CoV in Vero E6 cells (Russo et al., 2020). Investigating other potential anti-SARS-CoV-2 phytochemicals found in *A. annua* and *A. afra* is warranted.

## 5.0 Conclusions

Hot-water (tea infusion) extracts of *A. annua* are active against SARS-CoV-2 and its variants alpha, beta, gamma, delta, and kappa. In our original report, anti-SARS-CoV-2 activity inversely correlated with ART content. Herein, similar responses are noted for gamma, delta, and kappa wherein the *A. annua* cultivar with the lowest ART content, BUR, generally had the lowest (most effective) IC_50_. These results demonstrate the potential of the extracts as treatments in the global fight against this constantly evolving virus. We urge WHO to consider including extracts and encapsulated dried leaves in their announced clinical trials that already include artesunate (Kupferschmidt, 2021). We aim to test preclinical models of SARS-CoV-2 in rodent models (Dinnon et al., 2020; Gu et al., 2020) that could help advance *A. annua* as an inexpensive therapeutic in parts of the world where logistic issues such as delivery require longer time to achieve vaccination levels that would ultimately quell this pandemic.

## 6.0 Acknowledgements

We thank Prof. David Ho for supporting the live virus work in his lab. Award Number NIH-2R15AT008277-02 to PJW from the National Center for Complementary and Integrative Health funded phytochemical analyses of the plant material used in this study. The content is solely the responsibility of the authors and does not necessarily represent the official views of the National Center for Complementary and Integrative Health or the National Institutes of Health.

## 8.0 Author Contributions

Manoj Nair: Conceptualization; Data curation; Formal analysis; Investigation; Writing - review & editing.

Yaoxing Huang: Data curation; Formal analysis; Investigation;

David Fidock: Data curation; Resources; Writing - review & editing

Melissa Towler: Resources; Writing - review & editing.

Pamela Weathers: Conceptualization; Resources; Roles/Writing - original draft; Writing - review & editing.

